# Perception of energy production from biomass: A Mexican population pilot survey

**DOI:** 10.1101/2023.06.27.546801

**Authors:** J. A. Mascorro-Guzmán, J. Vásquez-Olmos, O. Zúñiga-Sánchez, E. Monteros-Curiel, L. C. Durand-Moreno, V. Flores-Payan, A. A. Angulo-Sherman

## Abstract

The term biomass may be used in different contexts, one of which is related to energy production from biomass sources, especially wastes. Mexico is considered a country with a high potential to exploit renewable energy sources, particularly biomass. Still, according to studies about energy consumption in Mexican homes, biomass is one of the least used, compared to conventional sources or even other renewable sources. It is not understood how aware the country’s population is of the potential of biomass, or if they recognize it as a renewable energy source. This study presents a pilot survey performed on a small adult population that has or is receiving professional education and designed to identify how biomass is perceived among other renewable energy sources and whether the population considers it relevant to inform themselves about this topic. The results obtained indicate that people recognize mostly wind (99%) and solar (88%) power, but when it comes to biomass only 36% of the population knows energy may be obtained from it. When it comes to its use only 5.8% of the population indicates using it. Besides understanding the population’s perception these results may help to develop more educational policies about renewable energies by universities.

## Introduction

The term biomass has two different interpretations, the first one, also the oldest refers to the quantity of matter that constitutes the living beings in a habitat, which is usually represented per area or volume; the second refers to the mass obtained from plants, animal wastes or any organic matter produced during a biological process, either spontaneously or in purpose, and that is used to as fuels [Merriam-Webster (2022), Real Academia Española (2022)]. Nevertheless, as mentioned in the Encyclopedia Britannica, depending on the area of knowledge in which it is used, we may find more specific definitions [Brittanica (2022)]. Then, it is possible that depending on the studies or experience of an individual, a certain meaning is accepted because of the frequency it is used to represent a particular idea.

In 2018, Mexico’s National Institute of Statistics and Geography (INEGI), conducted a survey with the purpose to obtain statistical information that allows knowing the consumption patterns of Mexican homes [SENER (2018)]. This survey is known as the National Survey on Energy Consumption in Private Homes (ENCEVI) and is presented by INEGI as the first survey applied to the Mexican population to obtain information on energy consumption in the different kinds of Mexican homes. According to the ENCEVI results, for the electric supply the homes that use renewable energy sources represent only 1%, from which the main source is through solar panels, specifically 0.25% of the homes use solely solar power, while the rest (0.75%), uses hybrid and bidirectional systems, that need to be connected to the public electricity supply network. Besides, for other domestic activities that require heating, solar energy is also the main renewable energy used: 28% of the houses use water solar heating systems. Although is not mentioned as biomass, among this survey results it is mentioned that some homes keep using firewood, which together with coal, are used in 11% of the homes for cooking activities, and in 10% for water heating.

There is a study almost contemporary to the ENCEVI survey, published in 2017 [Perez-Denicia E., et.al (2017)], that analyzed the exploitation of renewable sources for electric power production in Mexico. This work summarizes the status of electricity production from renewable sources in the country, as well as its energy potential. Among the variety of renewable sources, biomass is included and is expected that its energy potential is near 2396 GWh per year. In 2017, the focus of publications about the exploitation of renewable sources in Mexico could be distributed as follows; 41% about solar power, 26% related to biomass, 15% corresponding to wind power, and 10% for any other renewable sources [Perez-Denicia E., et.al (2017)]. This information makes evident the relevance that biomass exploitation should have, still, even including the public policies embraced by the country related to the use of renewable energies, the same seems to have no influence in promoting biomass utilization [Sanchez, S. F., Segovia, M. A. F., & López, L. C. R. (2023), Pischke, E. C., et.al (2019)]. Sanchez et al. mentioned that since the ‘70s up to 2020, the sources used for electric energy production in Mexico have shown a tendency to increase the consumption of non-renewable sources, reaching an amount of utilization of 82%, from which almost 72% is provided by thermoelectric plants, and 10% by coal [Sanchez, S. F., Segovia, M. A. F., & López, L. C. R. (2023)]. While for renewable sources the decrease may be exemplified using of hydroelectric plants which provided 57% of the energy in the ‘70s and its use decreased to 9% in 2020 [Sanchez, S. F., Segovia, M. A. F., & López, L. C. R. (2023)]. For the other renewable energies, it is mentioned that wind power has increased its use from 2015 to 2020, reaching a 6%, while the other sources, including biomass, are contained in the remaining 3%. In particular, we intend to highlight the potential use of biomass as an energy source, because when implemented correctly and considering the appropriate biomass treatment [Valdez-Vazquez, I., Acevedo-Benítez, J. A., & Hernández-Santiago, C. (2010)], for certain communities it may represent a more stable source [Sanchez, S. F., Segovia, M. A. F., & López, L. C. R. (2023)] compared to other renewable sources. Nevertheless, it is important to understand the limits of its utilization according to the way the national energy demand is increasing [Rios, M., & Kaltschmitt, M. (2013)].

To obtain energy from biomass there are three ways proposed: 1) heat and power, 2) biofuels and 3) biogas. The first refers to the traditional use of biomass for obtaining energy, like burning wood. The second is related to the production of liquid fuels like biodiesel and bioethanol, for this purpose, the biomass should receive either biochemical or thermochemical treatments to obtain the fuels. In the third one, biomass also is treated by anaerobic digestion, and the fuel resulting is a gas that includes methane along with other compounds [Perez-Denicia E., et.al (2017)]. All the resultant fuels are burned to obtain heat, which can be directly used or converted into electricity. It is suggested that the best way to obtain energy from biomass in Mexico is by burning, either biomass directly or biogas. Also, to obtain more benefits biomass should come from residues of crops other than sugar cane bagasse and firewood, which are the more commonly used in the country. Employing other biomass sources and providing a specific and appropriate treatment for its use, will allow cleaner and more secure exploitation for electric energy production, and consequently the promotion of biomass utilization in the Mexican energy basket [Perez-Denicia E., et.al (2017)].

Being aware of the potential use of biomass sources is important. On one side, it is believed that the methodologies and technologies used to transform biomass into electric energy have a long-term positive environmental impact because of the cost-benefit ratio. On the other side, the use of crops like jatropha, oil palm, corn, soy, or sugar cane, implies a higher environmental impact compared to the use of fossil fuels. The late is because the use of fossil fuels results in lower greenhouse gas emissions and the use of such crops is associated with high environmental and ecological costs, due to biodiversity loss, agricultural land degradation, and environmental pollution in general [Perez-Denicia E., et.al (2017)].

There is no doubt that Mexico has great energy potential by exploiting biomass sources. It is then relevant to identify whether the Mexican population is aware of the term biomass as an option among renewable energy sources, as well as recognize the existence of the resultant bioenergy and biofuels. In this work, we analyze the awareness and perspectives of these terms through a pilot survey performed on a group of Mexican adults.

## Methodology

This work analyzes the awareness of the term biomass as a renewable energy source. For this purpose, a pilot survey was applied to a group of 84 Mexican adults that are either studying or have studied at the university. This survey was designed using google forms and applied over a period of a month between October and November of 2022. The survey has 15 items, of which 3 are related to the professional area of study, 4 evaluate the previous knowledge of renewable sources, and 8 are associated with the term biomass, bioenergy, and the biofuels that may be obtained from this source.

To determine the survey reliability, its Cronbach’s alpha coefficient was calculated using equation 1; where k refers to the number of items, S_i_^2^ is the associated variance to the *i*th item, and S_t_^2^ is the associated variance of the total score, obtained by summing all the items [Ledesma, R., Molina, G., & Valero, P. (2002)].

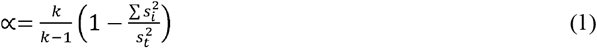

The calculation was done both, with Excel and with SPSS software. From the Excel results, considering all items, it was obtained a value □=0.68. It was determined that two items should not be considered. The first one asked about knowing the term renewable source, for which the whole population of the sample answered affirmatively. The second was a nominal qualitative item that applied only to the students in the population, to determine what semester was the student in. The information related to the second question eliminated is included in the results with the sole purpose to exemplify at what stage of their studies the students are.

Once it was determined these items would be excluded from the analysis the result for Cronbach’s alpha coefficient was □=0.848, which is considered a good reliability value [George, D., & Mallery, P. (2003)].

## Results

From the sample of 84 adults, it was determined that 64% of them are studying at the university, while 36% have already finished their studies. This sample included individuals from different professional areas of study, the most representative of those were Engineering and Technology (39%) and Natural and Exact sciences (32%), followed by Health sciences (13%), Social and Administrative sciences (11%) and Education, Humanities and Arts (5%). The student population is classified into three groups according to the semester they are in; from 1^st^ to 3^rd^ semester, it is said they are at the beginning-stage, from 4^th^ to 6^th^ semester they are classified as intermediate-stage students, from 7^th^ to 9^th^ semester they are considered as final-stage students. From the student population, 60.4% are considered students in the final-stage, 30.3% are students in the intermediate-stage, and 9.5% are beginning-stage students.

Every participant had previously heard about renewable energy sources. The most known sources are wind (100%), and solar (99%). These energy sources are followed by hydraulic (88%). Almost half of the people identify both geothermal and tidal energy. Finally, biomass is only recognized by 36% of the population. These results are presented in figure 1.

**Figure 1.**
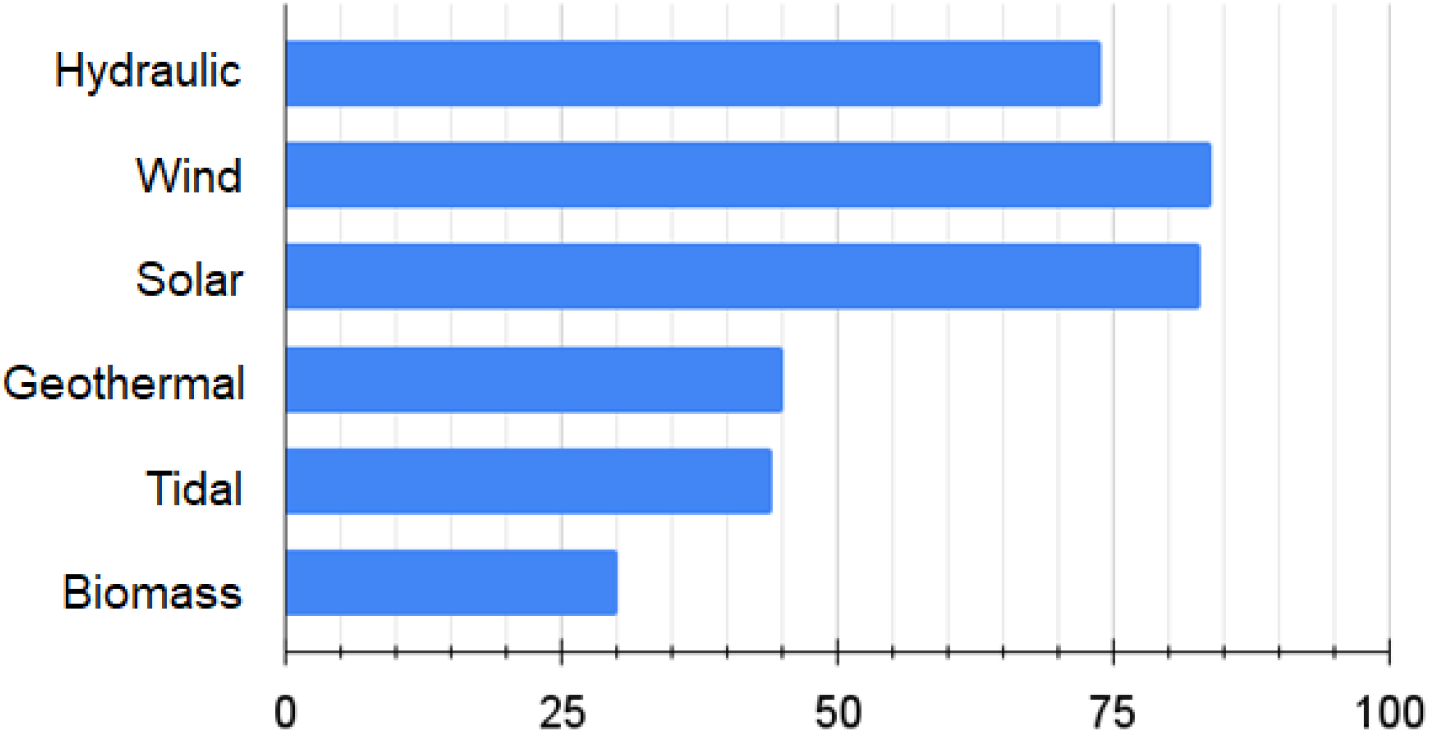
Frequency with which people have heard about the different renewable energies. The most recognized are wind and solar, followed by hydraulics. Half of the population knows about geothermal and tidal energy. Biomass energy is the least known.

Although the whole population has heard about at least one renewable energy source, only 62% use them daily. It is important to notice that this part of the population mentions using only one of the sources and not a combination of them. Also, from the six options provided, people only selected either biomass, hydraulic, solar, or wind. The one acknowledge as the most commonly used is solar energy, by 43 individuals, followed by hydraulic used by 5 of them, biomass used by 3, and only one person acknowledge to use of wind energy. The energy sources that the population is using are presented according to their percentages in figure 2.

**Figure 2.**
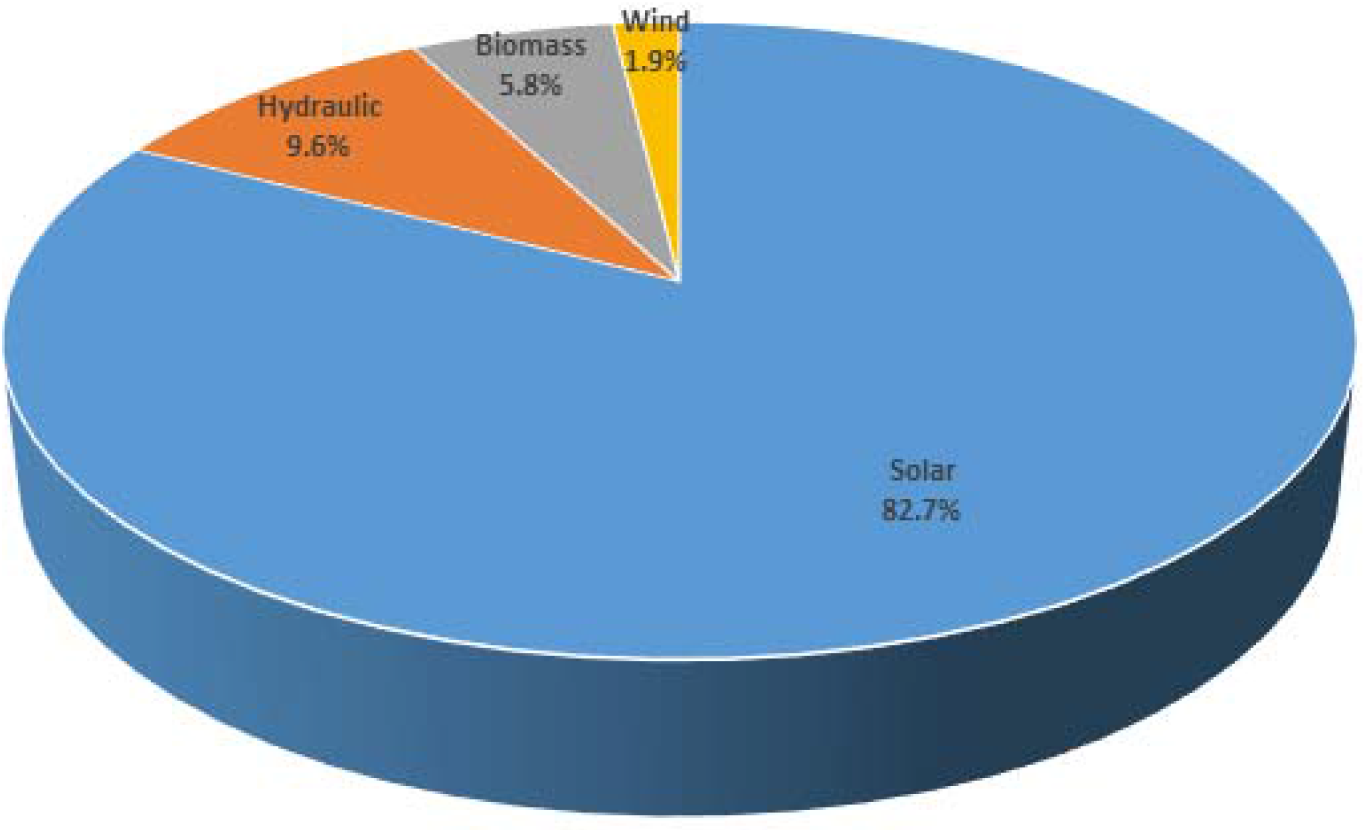
Energy sources that people who use renewable energy acknowledge using.

The results of the survey related to the terms biomass and bioenergy are the following: 57% of the population acknowledge hearing the term biomass, although only 36% relate it to an energy source. Besides, only 69% of the participants recognize that biomass and bioenergy are different terms, the rest believed they are the same. When asked about hearing previously of the term bioenergy, 79% answered affirmatively.

The participants in the survey were provided with four main classifications for biomass sources: agricultural, energy crop, natural, and residual, to identify if they recognize any. 42 individuals identified the term Residual biomass; the terms Natural biomass, and Agricultural biomass were recognized by 34 and 30 people respectively. The least popular classification was Energy crop, acknowledged only by 17 participants. These results are presented in figure 3A.

**Figure 3.**
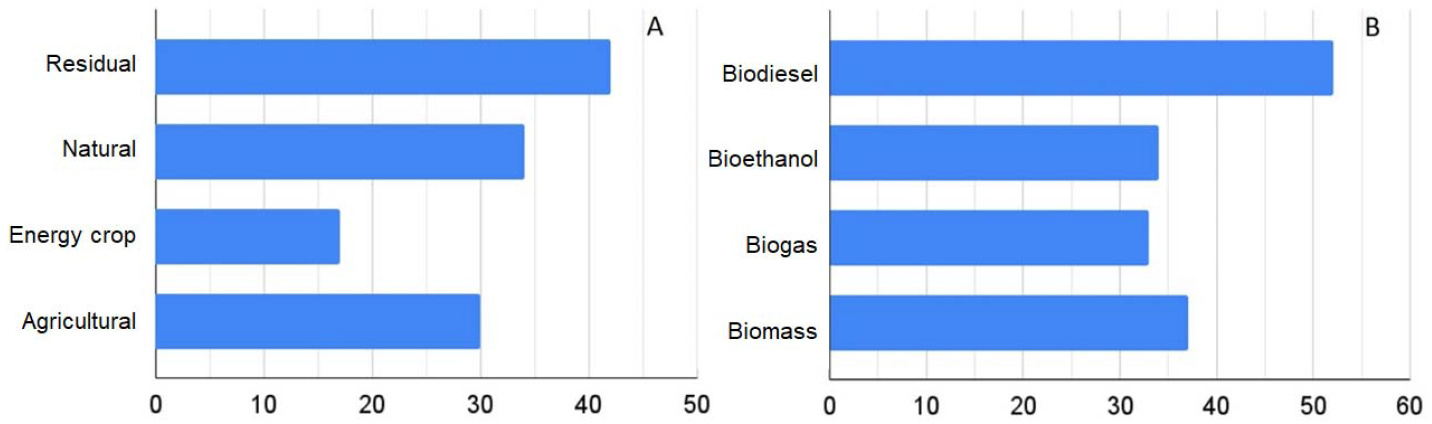
A) Kind of biomass that the sample population acknowledges, B) Biofuels that the sample population recognizes.

The survey then provided four bioenergy fuel options: biodiesel, bioethanol, biogas, and biomass. From these terms, the most known is biodiesel, recognized by 52, followed by biomass, bioethanol, and biogas, recognized by 37, 34, and 33 individuals respectively. These results appear in figure 3B.

Finally, information about the relevance of these terms and whether there is a negative impact on the environment by using biomass and biofuels to generate energy. According to the answers of the sample population, 99% find biomass and bioenergy useful, and 93% would like to receive more information about them. About whether the exploitation of biomass and biofuels may have a negative impact on the environment, only 13% believe so.

## Discussion

It is important to notice through the results that independently of the professional area of study, all individuals recognize renewable energy sources. Besides, the results obtained through this survey indicate that 83% of the population uses solar energy, which corresponds to the ENCEVI survey results, which indicated that of the Mexican houses that use these sources (1%), 25% use exclusively solar energy, while 75% come from hybrid systems; that use in addition of solar energy another complementary source. Also, it is important to notice from the results obtained in this work, that the most known sources are solar and wind (99%, and 100%), solar energy is widely used by the population in the survey that claims to use renewable energy, but when it comes to wind, the amount is only 1.9%.

ENCEVI survey already indicates that biomass, in the way of wood, is used in some Mexican homes as a fuel [SENER (2018)], but other biofuels are not considered, although the potential they may have as energy sources [Perez-Denicia E., et.al (2017)]. According to this work pilot survey, biomass is the least known renewable energy source, even though the academic community identifies it as Mexico’s second most interesting energy source. Keeping this in mind, it seems natural that according to this survey population, the utilization of biomass is least than 6%. On one hand, the use of biomass in Mexican homes is below 11% according to ENCEVI, which agrees with the results we have obtained, on the other hand, even though the academic community efforts to promote the use of biomass, it is evident that not all the survey population to identify its potential compared to other renewable energies which may explain why its exploitation is so scarce and is only used in its most rudimentary way as a fuel [SENER (2018),Perez-Denicia E., et.al (2017)].

Keeping in mind the previous ideas, it becomes relevant the type of biomass and biofuels the population identifies. The term residual biomass is the class of biomass most identified, while the least known is energy crop. The identification of the first term may be associated with the one of organic residues, which is a more common concept associated to residue recollection. In contrast, the recognition of the term energy crop, which corresponds to the production of crops exclusively for the fabrication of biofuels, becomes an example that in general, the population does not relate the term biomass with the generation of energy. If people are unable to relate these ideas, it is understandable why there may be least interest in using biomass compared to other renewable energy sources like those from wind or the sun.

Analyzing in particular the biofuels identified, it is important to compare the high recognition of the term biodiesel compared to the least known, biogas. This is because according to the academic community, the production of biofuels such as biodiesel or bioethanol is not the best option for this country to exploit biomass, and the efforts should be focused on either the production of solid biofuels, such as wood, or the production of biogas [Perez-Denicia E., et.al (2017)]. It is convenient to provide the Mexican population with information about the potential use of biomass and biofuels as an alternative energy source, expecting this increase may have an impact on public policies. It is encouraging that the population interviewed agreed almost totally that they found these topics relevant and even more that they would like to know more about them.

Finally, it is convenient to notice that most of the population believed that there isn’t a negative impact on the environment if these energy sources are exploited. The impact may depend on the process itself from the production of the raw material to the transformation into usable energy [Perez-Denicia E. et.al (2017)]. It is recommended that if biomass is used as an energy alternative, it must be done in a measured and responsible way.

## Conclusion

Mexico’s potential exploitation of biomass as a renewable energy source is promising, still, it appears that a significant part of the population is not used to relate this source to the production of energy.

One of the Universities’ substantive tasks must be the dissemination of science and knowledge, because of this, strategies to bring this knowledge about biomass and biofuels to the Mexican population should be encouraged. That way people may be able to promote the use of more energy sources alternatives, understanding both, the advantages of its use as well as the negative implications on the environment depending on the whole process of transforming it into usable energy.

This survey provides information that is relevant, pertinent, and reliable. Nevertheless, is convenient for it to be applied to a more extensive population, so a better strategy to enlighten people with this information in an accessible way may be designed.

## Author contributions

Angulo-Sherman A. A. proposed the idea and relevance of the realization of this survey, also designed the items, and promoted the survey through different digital public forums with the help of bachelor students Vazquez-Olmos J. and Mascorro-Guzmán J. A. Angulo-Sherman A. A. and Mascorro-Guzmán J. A. analyzed the results obtained through the survey. Zuñiga-Sanchez O. performed the reliability analysis of the survey through the SPSS software. Monteros-Curiel E. and Durand-Moreno L. C. provide meaningful feedback on the survey design and the document. Angulo-Sherman A. A. and Flores-Payan V. reviewed the final version of the document for its publication.

## Acknowledgments

The authors would like to thank bachelor student Alan Francisco Lopez Mijangos for his valuable contribution to the realization of this survey, and its promotion through different digital public forums. Also, Camacho-Rodriguez A., because of the significant feedback during the realization of this survey.

## Data availability

The data used to support the findings of this study are available from the corresponding author upon request.

